# *Eimeria falciformis* BayerHaberkorn1970 and novel wild derived isolates from house mice: differences in parasite lifecycle, pathogenicity and host immune reactions

**DOI:** 10.1101/611277

**Authors:** E. Al-khlifeh, A. Balard, V.H. Jarquín-Díaz, A. Weyrich, G. Wibbelt, E. Heitlinger

## Abstract

Species of *Eimeria* (Apicomplexa:Coccidia) differ in the timing of lifecycle progression and resulting infections vary in host immune reactions and pathology they induce. *Eimeria* infections in house mice are used as models for basic immunology and the most commonly used isolates have been passaged in laboratory mice for over 50 years. We questioned in how far such isolates are still representative for infections in natural systems.

In the current study, we address this question by comparing the “laboratory isolate” *E. falciformis* BayerHaberkorn1970 with a novel, wild derived isolate *E. falciformis* Brandenburg88, and contrast this with another novel wild derived isolate, *E. ferrisi* Brandenburg64. We compare parasite lifecycle progression. We relate this to immune cell infiltration at the site of infection (in the caecum) and cytokine gene expression in the spleen as a measure of host immune response. We assess host weight loss as a measure of pathogenicity.

A species-specific slower parasite lifecyle progression and higher pathogenicity are observed for *E. falciformis vs. E. ferrisi.* Host cytokines, in contrast, are expressed at significantly higher level in the spleen of mice infected with the *E. falciformis* laboratory isolate than in both wild derived isolates, irrespective of the species. Differences in histopathology are observable between all three isolates: The *E. falciformis* BayerHaberkorn1970 laboratory isolate induces the strongest inflammation and cellular infiltration (with lymphocytes, plasma cells and eosinophilic granulocytes) followed by the wild derived *E. falciformis* Brandenburg88 isolate. *E. ferrisi* Brandenburg64 is inducing milder histological changes than both *E. falciformis* isolates.

It can be speculated that the serial passaging of *E. falciformis* BayerHaberkorn1970 has resulted in evolutionary divergence rendering this isolate more virulent in NMRI mice. Caution is needed when findings from experimental infection with laboratory strains should be integrated with observations in natural systems.

**Highlights:**

- *E. ferrisi* has a shorter pre-patency than *w*ild-derived and laboratory isolates of *E. falciformis*.
- *E. ferrisi* is less virulent than both *E. falciformis* isolates and the timing of maximal oocyst shedding relative to host weight loss differs.
- The laboratory strain of *E. falciformis* induces stronger cytokine expression in the spleen than both wild derived strains of *E. falciformis* and *E. ferrisi*.
- The laboratory strain of *E. falciformis* induces stronger tissue infiltration of immune cells than the wild-derived strain. *E. ferrisi* infections are associated with the lowest infiltration.

## Introduction

Maintenance of parasite lifecycles via serial passaging is a cornerstone of experimental parasitology. Parasites are propagated under defined and controlled conditions with the aim to provide infective stages for experiments (Lucius et al., 2017). The procedure allows the parasite to evolve due to mutation and genetic drift or adaptation to the passaging host and environment (Ebert, 1998; Burke, 2012). Genetic drift is promoted by the use of small inocula during passaging, and drift can act while diversity of a parasite isolate is reduced intentionally to obtain a clonal strain. In clonal strains mutation and drift can continue to act (Farrell et al., 2014). Considering adaptive evolution, an important part of the passaging environment is given by living hosts (Elena and Lenski, 2003), which usually have low genetic diversity (e.g., clonal or inbred lines, cultures), are immunologically naive due to the absence of previous infections (Mackinnon and Read 2004) and lack co-infections with other parasites (Abolins et al., 2017). Procedures for serial passaging of parasites typically collect infective stages at a particular time after infection and use the obtained inoculum to infect new animals or use haphazard infections in dense environments. Iteration of such a static routine likely differs from natural parasite environments with different timing of infections and variable transmission. In most cases both the biotic (host) and abiotic (outer environment) during passaging differs profoundly from the environment experienced by the parasite during its life cycle under natural conditions. To summarize, parasite laboratory isolates might experience both neutral and adaptive evolutionary processes. As a consequence they might not be representative for analogues in natural systems.

Serial passage leads in most cases to higher virulence (enhanced growth and reproduction of the parasite, and larger impact on the host) in the host type used for the process (reviewed by Ebert, 1998). This can be due to low genetic diversity in host populations (for example inbred lines) used for passage, reducing fitness trade-offs associated with specialization and promoting the expansion of more virulent pathogens. This phenomenon has been demonstrated in systems including the apicomplexan parasites *Plasmodium* spp. in rodents (Mackinnon and Read, 1999, 2004; Barclay et al., 2014). Adaptation to the passage host in these studies increases parasite virulence.

Contrary, but still consistent with this, passage of highly virulent isolates of the apicomplexan parasite *Eimeria* spp. can lead to attenuation of virulence when only the first oocystes committing to sexual reproductions are selected (McDonald and Ballingall 1983; Shirley and Bellatti 1988; Matsubayashi et al., 2016). These attenuated strains are called “precocious lines” and are the basis for live vaccines used in the poultry industry (Shirley and Long, 1990). Given the practical implications of this phenomenon, numerous experiments focused on changes in parasite life history, virulence and host response that arise as a consequence of attenuation. It is clear from these experiments that *Eimeria* spp. respond quickly to selection pressure. In contrast – to our knowledge – no studies attempt to correlate enhanced virulence after serial passaging of *Eimeria* with physiological (e.g. immune-) responses in the passaging host.

Species of the genus *Eimeria* usually have a small host range, often infecting a single host species (Hnida and Duszynski, 1999; Kvičerová and Hypša, 2013; Hashimoto et al., 2014; Vrba and Pakandl, 2015) and reside at specific sites within the intestines of their hosts (Haberkorn, 1970; Owen, 1975; Chapman et al., 2013). All species have a direct life cycle with asexual expansion and sexual reproduction within epithelial cells of the gastrointestinal tract before transmission stages (oocysts) are released. Diploid oocysts become infective after reductive divisions (sporulation) in the environment (Canning and Anwar, 1968).

*Eimeria* spp. are widespread in diverse host species including many vertebrates from mammals to fish (Molnár et al., 2012). Infection causes damage in the intestinal mucosa resulting in malabsorption of nutrients and weight loss (Haberkorn, 1970; Chapman et al., 2013). As this has an economic impact in livestock, coccidiosis is an important focus in veterinary research (Brake et al., 1997; Laurent et al., 2001; Gadde et al., 2009; Swaggerty et al., 2011). *Eimeria* species capable of natural infection of the house mouse (*Mus musculus*) have been proposed as a model for e.g. host immune response against *Eimeria* (Heitlinger et al., 2014; Schmid et al., 2014). Serial passaging of laboratory isolates of *Eimeria* is usually conducted by collecting oocysts at the day of peak shedding. Oocysts are sporulated in an aqueous solution of potassium dichromate and inocula are used for new infections two to six months later, before infectivity decreases. The isolate *E. falciformis* BayerHaberkorn1970 has been isolated in 1960 (Haberkorn 1970) and since has been propagated in laboratories (first at Bayer animal health, Monheim, Germany; then at the institute for molecular parasitology of the Humboldt University, Berlin, Germany). In nearly 60 years since its isolation *E. falciformis* BayerHaberkorn1970 has likely become the most commonly used laboratory isolate of rodent *Eimeria* (Mahrt and Shi 1988; Schito et al., 1996; Steinfelder et al., 2005; Pogonka et al., 2010; Stange et al., 2012; Schmid et al., 2014, 2012; Ehret et al., 2017).

In the present study we compared infection of mice (NMRI) with the laboratory isolate *E. falciformis* BayerHaberkorn1970, to infections with wild derived isolates of *E. falciformis* (novel isolate Brandenburg88) and *E. ferrisi* (Levine and Evens, 1965) (novel isolate Brandenburg64). We assessed similarities and differences in proliferation of tissue stages, oocyst shedding and in the host pathological changes and immune response between the three different *Eimeria* isolates.

## RESULTS

### Dynamics of infection and host body weight loss differ between *Eimeria* species

Genotyping showed that two novel isolates which we obtained from individual house mice captured in the federal state of Brandenburg (Germany) belong to the species *E. falciformis* and *E. ferrisi*. We compared fragments of DNA sequences of the mitochondrial Cytochrom C oxidase subunit I (COI), the nuclear small ribosomal subunit (18S) and the open reading frame 470 (ORF470) of the apicoplast genome. For all makers the Brandenburg88 and Brandenburg64 isolate showed 99-100% sequence identity to available sequences from *E. falciformis* and *E. ferrisi*, respectively. This identifies the isolates robustly as the respective species as described before (Jarquín-Díaz et al., 2019). We named the novel isolates *E. falciformis* Brandenburg88 and *E. ferrisi* Brandenburg64 after the area (Brandenburg) from which they were isolated and a running number for mice used in our sampling (mouse no. 64 and 88).

We infected NMRI mice with these novel *Eimeria* isolates as well as with the laboratory isolate *E. falciformis* BayerHaberkorn1970 and followed the progression of infection by measuring parasite reproduction and host body weight loss. We assessed parasite reproduction via oocyst shedding from two to eleven days post infection (dpi) (Figure 1a). The two *E. falciformis* isolates and *E. ferrisi* showed different infection dynamics in NMRI mice: oocyst shedding of *E. ferrisi* has its peak intensity at 6 dpi, was drastically reduced at 7 dpi (n = 12, U = 2.91, p = 0.002), and fell below detection levels at 10 dpi. Oocyst shedding of *E. falciformis* has a peak intensity at 8 dpi for the laboratory isolate BayerHaberkorn1970 and at 9 dpi for the novel isolate Brandenburg88. The oocyst numbers declined after this peak in both isolates, but shedding was still detectable at 11 dpi when we sacrificed the last mice. For the two *E. falciformis* isolates we observed no difference in shedding intensity of oocysts at the peak day (n = 12, U = 0.24, p = 0.846) and peak oocyst abundance did not differ significantly between *E. ferrisi* and both *E. falciformis* strains (*E. ferrisi vs. E. falciformis* Brandenburg88, n = 12, U = 0.32, p= 0.777; *E. ferrisi vs. E. falciformis* BayerHaberkorn1970, n = 12, U = 0.96, p= 0.37).

**Figure 1.**
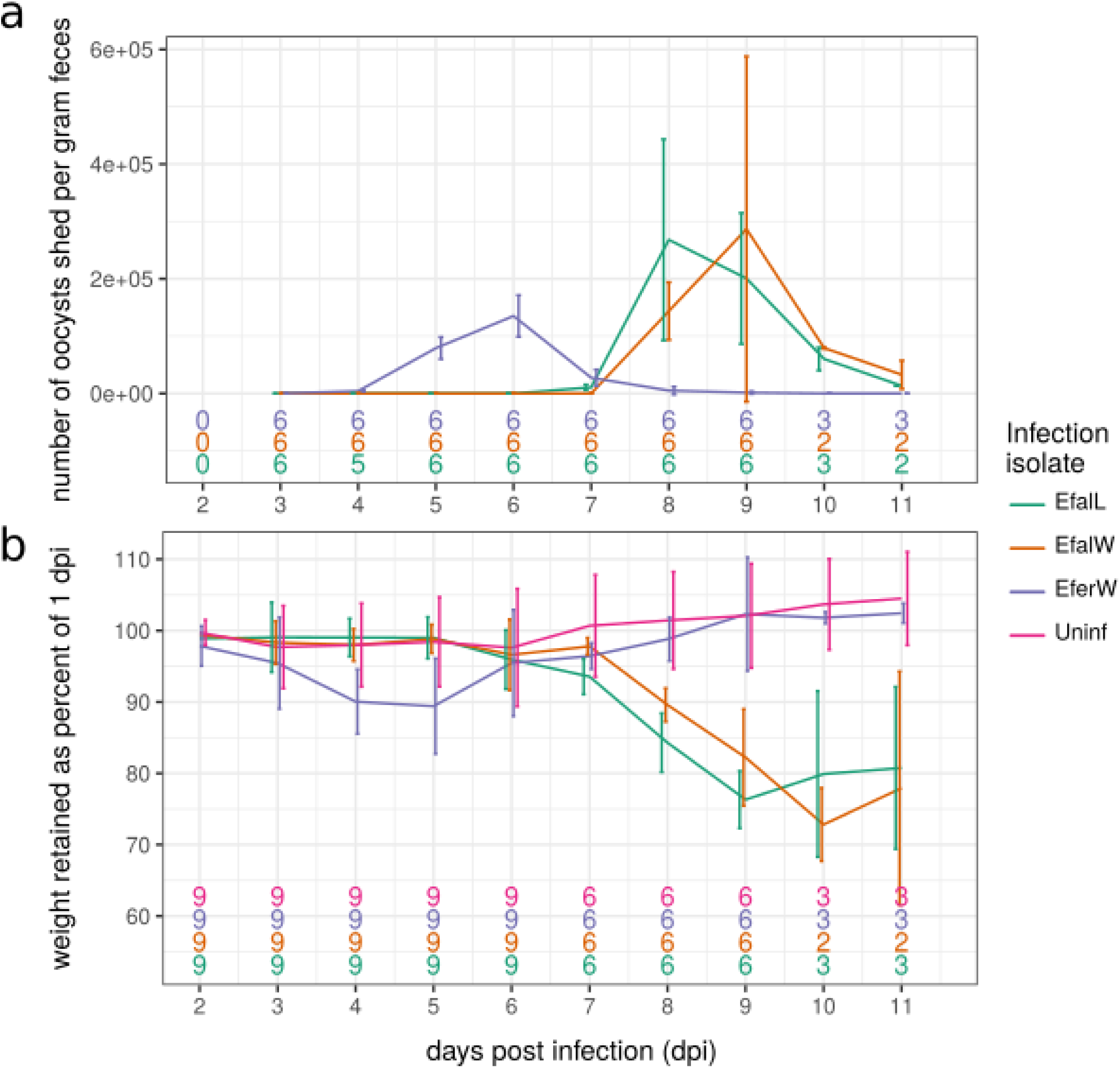
Parasite life cycle progression and pathogenicity assessed as host weight loss during the infection. Dynamics of parasite reproduction and host weight loss differ between the species *E. falciformis* and *E. ferrisi*. a) Oocyst shedding of *Eimeria* spp. from experimentally infected mice (NMRI) is displayed from 1 to 11 days post infection (dpi). Mice were infected with 200 sporulated oocyst of *E. ferrisi* Brandenburg64 (a recently derived isolate; EferW), *E. falciformis* BayerHaberkorn1970 (a classical laboratory isolate; EfalL) or and *E. falciformis* Brandenburg88 (a recently derived isolate, EfalW). b) Host body weight loss of the same three groups of mice is depicted as percentage of body weight retained compared to 1 dpi. The number of mice per group is given at the bottom of the plot. The number is gradually reduced towards the end of the experiment, because mice were sacrificed for collection of tissue samples. Lines indicate the mean for each group, error bars give the standard deviation.

The period of patency (oocyst shedding) was characterized by body weight loss in infected mice in all infections (Figure 1b). Infections with *E. ferrisi* coincided with significant weight loss at 4 dpi (n = 18, U = −2.43, p = 0.013) and 5 dpi (n = 18, U = −2.52, p = 0.010) in comparison to the control group. Infection with *E. falciformis* was accompanied by significant weight loss at 8 and 9 dpi in both *E. falciformis* BayerHaberkorn1970 (both dpi, n = 12, U = −2.89, p = 0.002) and *E. falciformis* Brandenburg88 (8dpi, n = 12, U = −2.41, p = 0.013; 9dpi, n = 12, U = −2.89, p = 0.002) isolates as compared to the control group. Weight losses in infections with *E. ferrisi* at their maximum (at 5 dpi) were, however, significantly lower compared to weight loss in infections with *E. falciformis* at their maximum (9 dpi; *E. ferrisi* vs. *E. falciformis* Brandenburg88, n = 15, U = −2.0, p = 0.049; *E. ferrisi* vs. *E. falciformis* BayerHaberkorn1970, n = 15, U = −2.59, p = 0.007).

Oocyst shedding and weight loss show different relative timing in *E. falciformis* compared to *E. ferrisi*. In infections with both isolates of *E. falciformis* weight loss coincides with or follows up to two days after oocyst shedding. In infections with *E. ferrisi* weight loss precedes peak oocyst shedding by one day or more (Figure 2).

**Figure 2.**
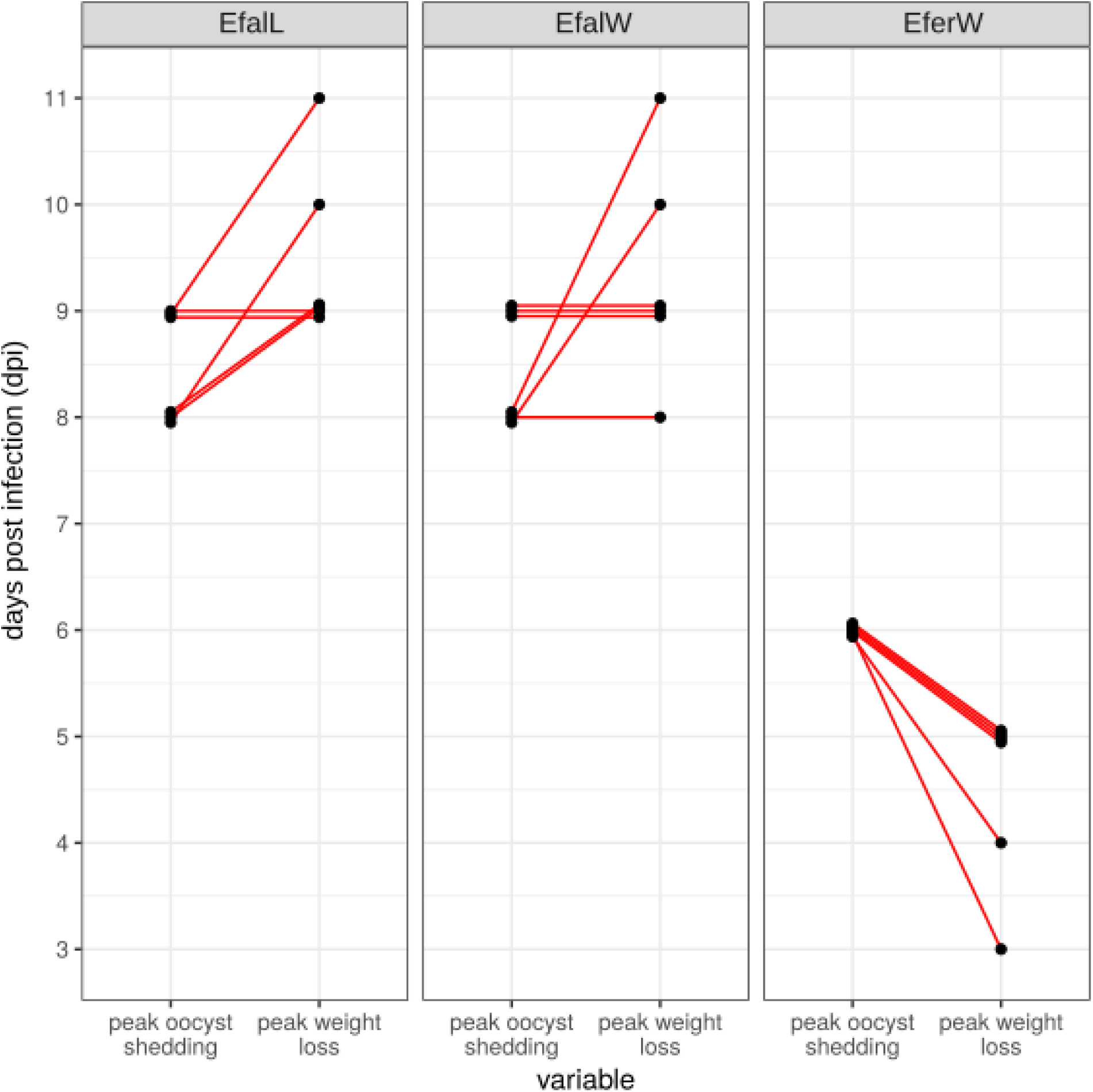
Relative timing of host weight loss and parasite reproduction differ between the species *E. falciformis* and *E. ferrisi* for individual hosts. The peak of the host’s weight loss follows after the peak of oocyst shedding in infections with *E. falciformis*, while in infections with *E. ferrisi* the hosts lost most weight before most parasite oocysts were shed. Points depict the peak day of both oocyst shedding and weight loss, respectively, red lines connect both measurement from the same mouse. All mice sacrificed at 9dpi or later are assessed. For underlying experimental procedures see Figure 1 and the Methods section. In case of *E. falciformis* infections the day of maximal weight loss could have been potentially even later if some mice (n = 3 for each isolate) were not sacrificed at 9 dpi.

### Intensity of tissue stages of *Eimeria spp*

*Eimeria* infections in our study result in a transient presence of parasite stages in epithelial cells of the caecum. We quantified the intensity of infection by a quantitative PCR (qPCR) assay using gDNA from caecal tissue. By specifically amplifying genes for the parasite (Cytochrome C-oxidase subunit I; COI) and the host (nuclear *cdc42* gene), we analysed the ratio of parasite DNA to host DNA. We report this ratio on a native (log2) scale of measurement and further refer to it as host-parasite ΔCt (Figure 3). The analysis of infected (*E. ferrisi* n = 15; *E. falciformis* Brandenburg88 n = 14; *E. falciformis* BayerHaberkorn1970 n = 14) and control samples (n = 13) allows us to estimate a limit of detection (LOD; mean + 2 standard deviations of the negative controls) for the assay at a host-parasite ΔCt of −3.73. This corresponds to eight *Eimeria* COI molecules for 100 copies of the mouse nuclear genome. We measured the highest value for an individual negative control sample at a host-parasite ΔCt of −4.84. Maximum values for host-parasite ΔCt (observed for the *E. falciformis* BayerHaberkorn1970 isolate) were 7.74, indicating a ratio of 214 parasite COI mDNA copies for each copy of the mouse genome in crude caecum tissue at this point.

**Figure 3.**
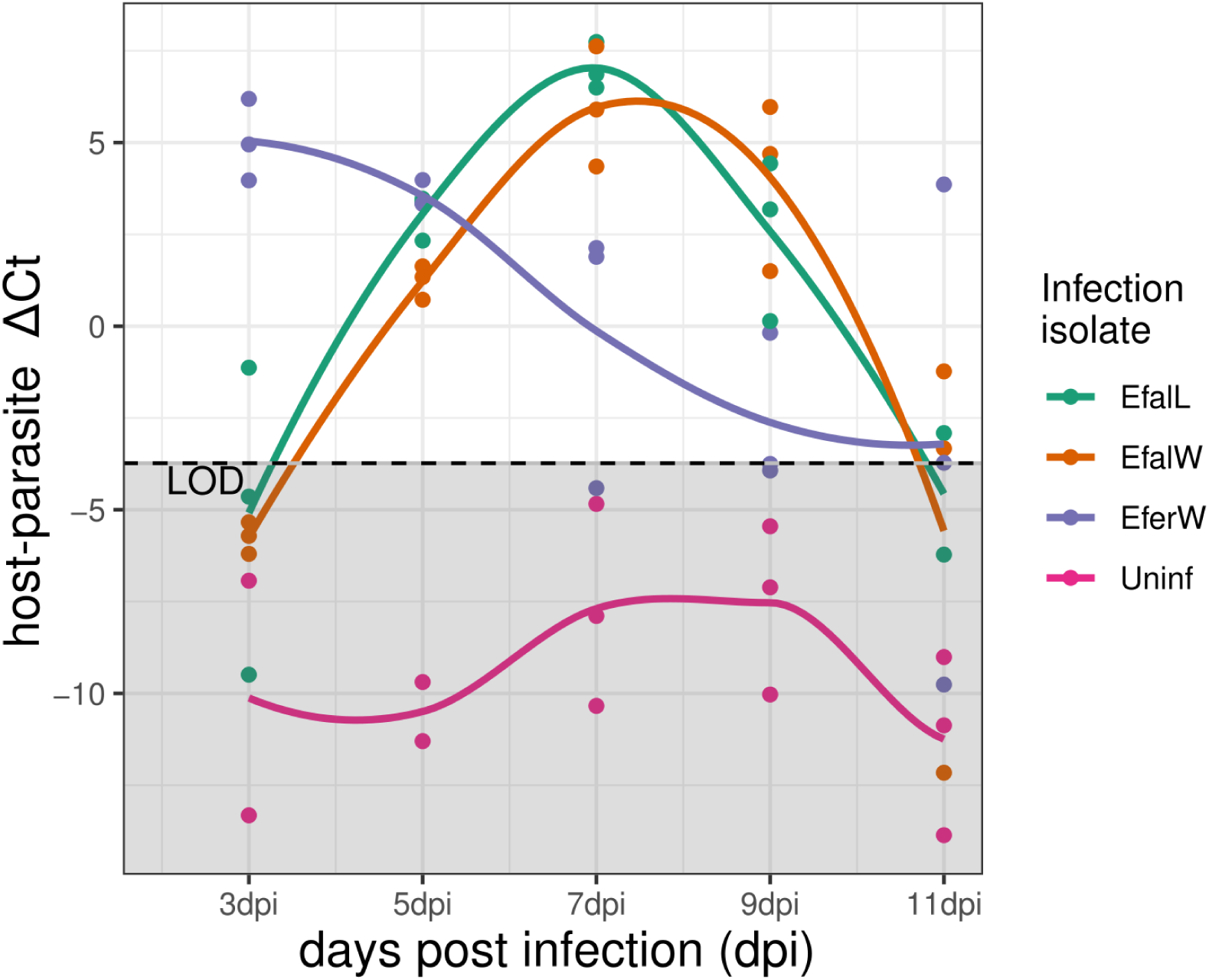
The relative amount of parasite *vs.* host DNA (host-parasite ΔCt) estimates the intensity of parasite stages in caecum tissue. The host-parasite ΔCt was calculated from the difference in cycle of threshold (Ct) values of qPCRs performed on a single copy nuclear gene (Cdc42) of the host and on a mitochondrial (COI) gene of the parasite. A limit of detection (LOD) was calculated as the mean + 2 standard deviations of ΔCt values of the negative controls. Coloured lines are drawn using local polynomial regression fitting (a “loess smoother”).

At 3 dpi *E. ferrisi* has the highest value of host-parasite ΔCt (6.19), while most infections with *E. falciformis* were still below the limit of detection (all *E. falciformis* Brandenburg88 and for two out of three *E. falciformis* BayerHaberkorn1970). For *E. falciformis* (both isolates) host-parasite ΔCt increased to values well above zero (equal numbers of parasite mitochondrial and host nuclear DNA copies) at 5 dpi and highest values were reached at 7 dpi. Again, the amount of DNA measured for *E. falciformis* (at this peak intensity) was similar to that of *E. ferrisi* (at 3 dpi, its peak). At 11 dpi the parasite-mouse ΔCt was reduced to values below zero for all samples, and for most samples below the limit of detection (except for one *E. ferrisi* sample, for which a value of 3.86 was measured).

### Differences in immune gene expression between the laboratory and wild-derived isolates

To characterize potential differences in the immune response of NMRI mice against the *Eimeria* isolates we assessed gene expression of relevant cytokines in the spleen. Expression levels for most genes differ significantly between uninfected controls and mice infected with the laboratory isolate *E. falciformis* BayerHaberkorn1970 (Figure 4). We used linear mixed effect models with dpi as random effect to “pool” information over multiple dpi, effectively increasing sample sizes (Table 1). Mice infected with *E. falciformis* BayerHaberkorn1970 show significantly higher expression levels of chemokine 9 (*CxCl9*), interleukins 10 and 12 (*Il10* and *Il12*), tumour growth factor beta (*Tgfβ*), and signal transducer and activator of transcription 6 (*Stat6*). We did not detect significant expression differences between control and *E. falciformis* BayerHaberkorn1970 infected mice for interleukin 6 (*Il6*) and interferon gamma (*Ifnγ*).

**Figure 4.**
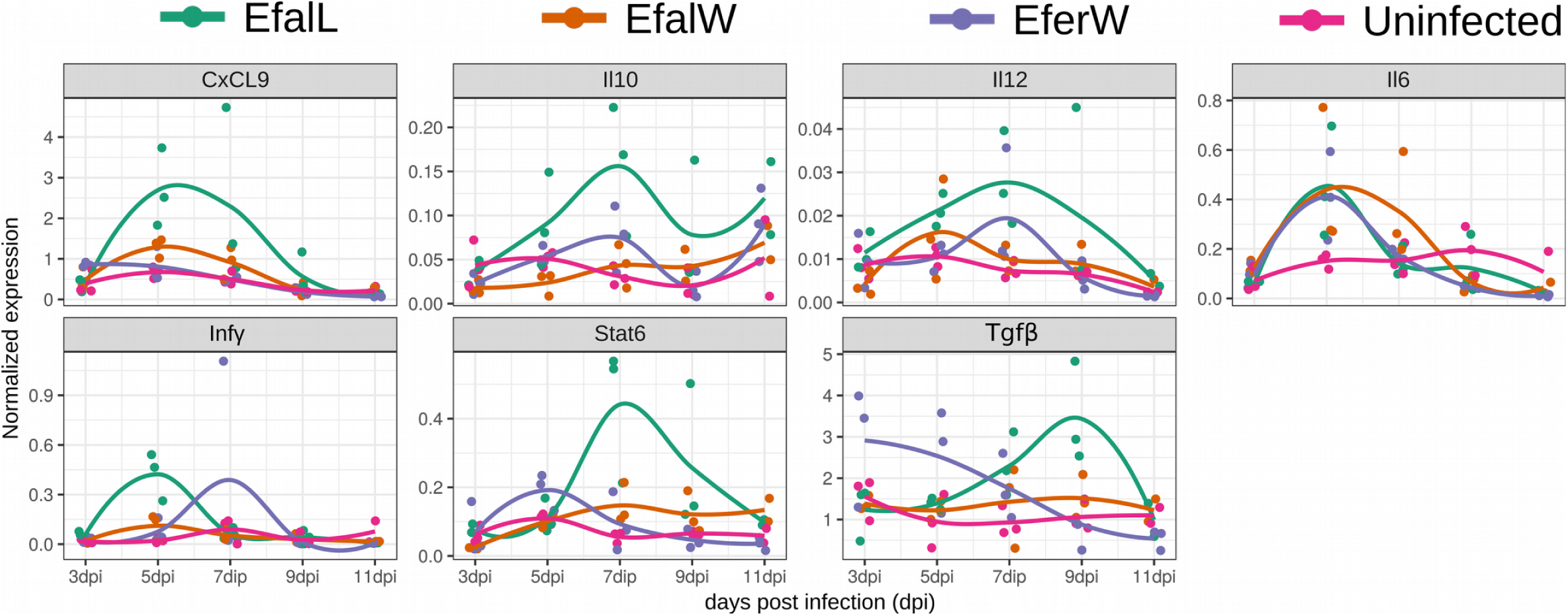
Mice infected with a laboratory isolate of *E. falciformis* showed elevated gene expression in the spleen. Mice (NMRI) were infected with the wild derived isolate *E. falciformis* Brandenburg88 (EfalW), the laboratory isolate *E. falciformis* BayerHaberkorn1970 (EfalL), and the wild derived isolate *E. ferrisi* Brandenburg64 (EferW). Only *E. falciformis* BayerHaberkorn1970 induces mRNA expression in the spleen elevated over non-infected control and over mice infected with both wild derived isolates. Mice were sacrificed at different time points post infection (dpi) and mRNA expression levels were assessed using quantitative PCR. Dots indicate normalized expression values for individual mice. Lines are drawn using local polynomial regression fitting (a “loess smoother”).

**Table 1:** Generalized linear mixed effect models show gene expression differences between wild (Brandenburg88; EfalW) and laboratory (BayerHaberkorn1970; EfalL) isolates of *E. falciformis*, a wild-derived isolate of *E. ferrisi* (Brandenburg64) and uninfected controls (Uninf).

|  | *CxCL9* |  |  | *Il10* |  |  | *Il12* |  |  | *Il6* |  |  | *Infy* |  |  | *Stat6* |  |  | *Tgfb* |  |  |
| --- | --- | --- | --- | --- | --- | --- | --- | --- | --- | --- | --- | --- | --- | --- | --- | --- | --- | --- | --- | --- | --- |
|  | *B* | *CI* | *p* | *B* | *CI* | *p* | *B* | *CI* | *p* | *B* | *CI* | *p* | *B* | *CI* | *p* | *B* | *CI* | *p* | *B* | *CI* | *p* |
| **Fixed Parts: Contrast against uninfected (intercept)** |  |  |  |  |  |  |  |  |  |  |  |  |  |  |  |  |  |  |  |  |  |
| (Intercept) | 0.39 | -0.15–0.92 | .194 | 0.04 | 0.01–0.07 | .012 | 0.01 | 0.00–0.01 | .035 | 0.13 | 0.00–0.26 | .087 | 0.04 | -0.05–0.14 | .400 | 0.07 | 0.01–0.13 | .045 | 1.11 | 0.61–1.60 | <.001 |
| inf.strain (EfalL) | 0.87 | 0.38–1.36 | **.001** | 0.06 | 0.03–0.08 | <**.001** | 0.01 | 0.00–0.02 | <**.001** | 0.03 | -0.06–0.13 | .499 | 0.08 | -0.04–0.20 | .194 | 0.13 | 0.06–0.21 | <**.001** | 0.81 | 0.14–1.48 | **.021** |
| inf.strain (EfalW) | 0.24 | -0.25–0.73 | .334 | -0.00 | -0.03–0.03 | .918 | 0.00 | -0.00–0.01 | .606 | 0.08 | -0.02–0.17 | .109 | 0.00 | -0.12–0.13 | .955 | 0.03 | -0.04–0.11 | .380 | 0.24 | -0.43–0.91 | .480 |
| inf.strain (EferW) | 0.10 | -0.38–0.58 | .694 | 0.01 | -0.01–0.04 | .374 | 0.00 | -0.00–0.01 | .403 | 0.02 | -0.08–0.11 | .721 | 0.06 | -0.06–0.18 | .353 | 0.02 | -0.05–0.09 | .622 | 0.61 | -0.04–1.27 | .074 |
| **Random Parts** |  |  |  |  |  |  |  |  |  |  |  |  |  |  |  |  |  |  |  |  |  |
| $\sigma^2$ | | 0.437 | | | 0.001 | | | 0.000 | | | 0.016 | | | 0.027 | | | 0.010 | | | 0.814 | |
| $\tau_{00, dpi.diss}$ | | 0.216 | | | 0.000 | | | 0.000 | | | 0.015 | | | 0.003 | | | 0.002 | | | 0.028 | |
| $N_{dpi.diss}$ | | 5 | | | 5 | | | 5 | | | 5 | | | 5 | | | 5 | | | 5 | |
| $ICC_{dpi.diss}$ | | 0.331 | | | 0.227 | | | 0.270 | | | 0.478 | | | 0.084 | | | 0.140 | | | 0.033 | |
| Observations |  | 57 |  |  | 57 |  |  | 57 |  |  | 57 |  |  | 57 |  |  | 57 |  |  | 57 |  |
| $R^2 / \Omega_0^2$ | | .456 / .453 | | | .440 / .437 | | | .421 / .417 | | | .498 / .495 | | | .162 / .147 | | | .342 / .338 | | | .157 / .154 | |
| **Fixed Parts: Contrast against *E. falciformis* BayerHaberkorn 1970 (intercept; same random effects)** |  |  |  |  |  |  |  |  |  |  |  |  |  |  |  |  |  |  |  |  |  |
| (Intercept) | 1.26 | 0.72–1.79 | .002 | 0.10 | 0.07–0.12 | <.001 | 0.02 | 0.01–0.02 | <.001 | 0.16 | 0.04–0.29 | .042 | 0.12 | 0.03–0.22 | .020 | 0.20 | 0.14–0.26 | <.001 | 1.92 | 1.43–2.42 | <.001 |
| inf.strain (EfalW) | -0.63 | -1.12–-0.14 | **.015** | -0.06 | -0.08–-0.03 | <**.001** | -0.01 | -0.01–-0.00 | **.003** | 0.05 | -0.05–0.14 | .347 | -0.08 | -0.20–0.04 | .213 | -0.10 | -0.17–-0.03 | **.010** | -0.57 | -1.24–0.10 | .101 |
| inf.strain (EferW) | -0.78 | -1.26–-0.29 | **.003** | -0.04 | -0.07–-0.02 | **.002** | -0.01 | -0.01–-0.00 | **.006** | -0.02 | -0.11–0.08 | .741 | -0.02 | -0.15–0.10 | .690 | -0.12 | -0.19–-0.04 | **.003** | -0.20 | -0.86–0.46 | .553 |
| inf.strain (Uninf) | -0.87 | -1.36–-0.38 | **.001** | -0.06 | -0.08–-0.03 | <**.001** | -0.01 | -0.02–-0.00 | <**.001** | -0.03 | -0.13–0.06 | .499 | -0.08 | -0.20–0.04 | .194 | -0.13 | -0.21–-0.06 | <**.001** | -0.81 | -1.48–-0.14 | .021 |

In contrast, for both wild-derived strains, *E. falciformis* Brandenburg88 and *E. ferrisi* Brandenburg64, expression levels for any of the examined genes do not differ significantly between uninfected and infected mice. Expression levels in infections with the laboratory isolate are significantly elevated also compared to infections with both wild derived parasite isolates (Table 1).

Some genes show (according to the model outlined above) non-significant differences in gene expression profiles over the negative control for the whole course of infection. This includes differences between infections with different parasite isolates (Figure 4). We did not analyse these differences on individual days statistically due to the low sample sizes, but give a description of our observations. *Il6* shows elevated levels of expression for all infection groups compared to controls at 5 dpi. Expression levels for *Ifnγ* seem elevated only at 5 dpi and only in infections with the *E. falciformis* BayerHaberkorn1970. Both cases of potential elevations in expression fail to be detected as significant over controls in our mixed effect models, likely because they are transient and diminished already at 7 dpi. *Il10, Il12, Stat6* and *CxCl9* show elevated expression levels at multiple days of infection with *E. falciformis* BayerHaberkorn1970 compared to all other infection groups (and were thus significant in our model). *Tgfβ* shows elevated expression levels early in infection with *E. ferrisi* (3 and 5 dpi) and late in infections with *E. falciformis* BayerHaberkorn (7 and 9 dpi). Taken together these observations add detail on the individual cytokines and underline our general finding of differences between wild-derived and laboratory isolates of *E. falciformis*.

In summary host gene expression of genes relevant for immune responses does not differ significantly from uninfected controls during infection with wild derived isolates of both *E. falciformis* and *E. ferrisi*. In contrast, most genes are expressed at significantly higher levels in infections with the laboratory isolate of *E. falciformis* BayerHaberkorn1970 compared to uninfected controls but also to all other infections including those with the wild derived *E. falciformis* Brandenburg88 isolate.

### Inflammatory cell infiltration differs between *Eimeria* isolates

To link our observation of gene expression in the spleen with independent measures of immune response and pathological changes, we performed a histological scoring of inflammatory cell infiltration (Table 2, Figure 5). Uninfected mice did not show inflammatory cell infiltration, besides a few (n = 2) exceptions with very low numbers of infiltrating lymphocytes.

**Table 2:** Score for the relative severity of leukocyte infiltration in histologic sections from the mid-part of the caecum from NMRI mice infected with *Eimeria* spp.

| Infection | Relative severity of leukocyte infiltration in caecum<sup>1</sup> |  |  |  |  |
| --- | --- | --- | --- | --- | --- |
|  | 3dpi | 5dpi | 7dpi | 9dpi | 11dpi |
| ***E. ferrisi* Brandenburg64** | 2, 1, 1 | 1, 2, 2 | 1, 2, 1 | 1, 1, 1 | 0, 0, 0 |
| ***E. falciformis* Brandnburg88** | 0, 1, 1 | 1, 0, 1 | 2, 2, 2 | 3, 2, 3 | 2, 2 |
| ***E. falciformis* BayerHaberKorn1970** | 1, 2, 1 | 3, 3, 3 | 3, 3, 2 | 3, 3, 2 | 1, 1 |

<sup>1</sup>Leukocyte infiltration was scored on a 0 to 3 scale, where 0 represent no infiltration and 1, 2, 3 represented low, moderate, or high infiltration, respectively. One section from each caecum sample was used for scoring. Three fields of view (at 200-time magnification) were evaluated for the amount of inflammatory infiltrates and a numerical score was assigned by averaging over these fields.

**Figure 5.**
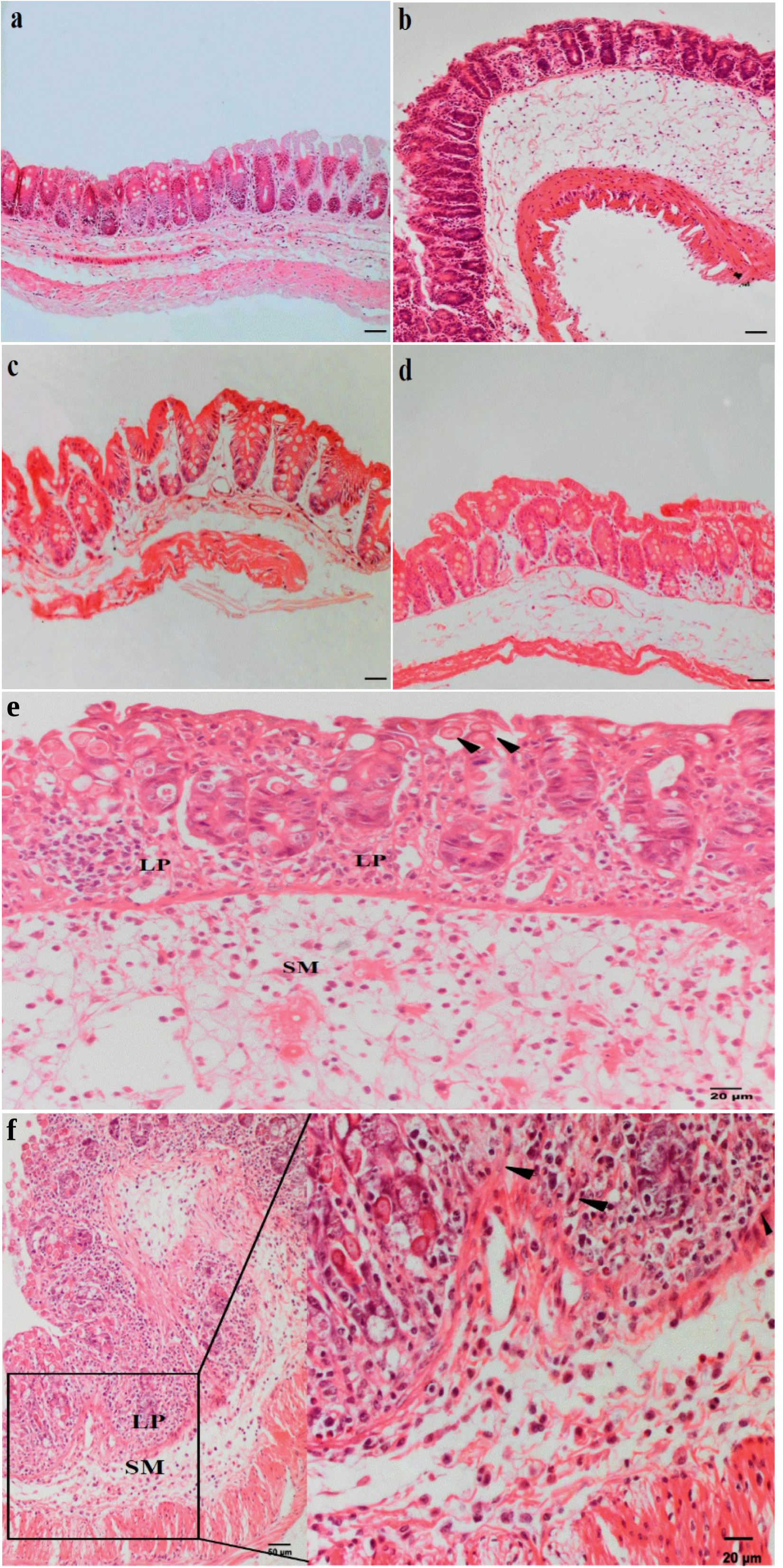
Immune cell infiltration and parasites in histological caecum sections obtained from mice infected with different *Eimeria* isolates. NMRI mice were innoculated with 200 oocysts. a) Mice infected with the laboratory strain *E. falciformis* BayerHaberkorn1970 show moderate inflammation (score 2, Table 2) at 3 dpi and strong inflammation b) at 5 dpi (score 3, Table 2). Infections with the wild derived isolate *E. ferrisi* Brandenburg64 at c) 3 dpi and d) 5 dpi, show low numbers of infiltrating immune cells (score 1, Table 2). Evident inflammatory process (score 3= high, Table 2) is associated with appearance of sexual stages and developing oocysts (black arrows) of the *E. falciformis* BayerHaberkorn1970 isolate at 7 dpi (e) and the wild derived isolate *E. falciformis* Brandenburg88 at 9 dpi (f). The infiltrate consists mainly of lymphocytes and plasma cells. Note the high cellularity in the mucosa comprising lymphocytes in the lamina propria (LP) with fewer eosinophil and neutrophile in the LP and submucosa (SM). Scale bar 50μm in a-d.

In mice infected with the laboratory isolate *E. falciformis* BayerHaberkorn1970 a relatively high score of inflammation was observed during pre-patency as early as at 5 dpi. The extent of immune infiltration remained high until 9 dpi and declined towards 11 dpi. In contrast, in caeca of mice infected with the wild derived isolate *E. falciformis* Brandenburg 88, only low numbers of inflammatory cells were found during the pre-patent period. Infiltration consisted mainly of lymphocytes and plasma cells at this stage of infection. Inflammation then increased at 7 dpi, and during this period, eosinophilic granulocytes were also detected. Infiltration was strongest at 9 dpi in infections with this isolate before decreasing slightly towards 11 dpi. The quality of the infiltration was the same in infection with both wild derived and laboratory isolate of *E. falciformis*. Infiltrations are consistently characterised by lymphocytes, plasma cells and the presence of eosinophils (Figure 5).

In infections with *E. ferrisi* Brandenburg64 a milder inflammatory response was detected on 3 dpi and 5 dpi, with the latter being relatively stronger (compared to other dpi of *E. ferrisi* infection), followed by a subsequent decline towards 7 dpi.

We analysed the inflammation score of mice sacrificed at different dpi (Table 2) in mixed effect models using dpi as random factor. Overall inflammation was significantly lower in *E. ferresi* infected mice than in those infected with *E. falciformis* (glmm; *E. ferrisi vs. E. falciformis* BayerHaberkorn1970, p = 0.001; *E. ferrisi vs. E. falciformis* Brandenburg88, p = 0.014).

## Discussion

Infections are increasingly studied in combinations of natural systems and laboratory settings to understand the evolution of pathogens and the immune system. Natural parasites of the house mouse can play an important role when bridging this research with experimental immunology. We here study *Eimeria* spp., natural parasites of the house mouse. We evaluated whether the “laboratory isolate” *E. falciformis* BayerHaberkorn1970 (Haberkorn, 1970) differs from wild derived isolates of the same species and of *E. ferrisi* concerning infection dynamics, pathogenicity and the immune reactions it induces. We found not only expected differences in parasite life cycle progression between *Eimeria* species (*E. falciformis* vs. *E. ferrisi*), but also differences between the wild derived and laboratory isolate of *E. falciformis.* The laboratory isolate of *E. falciformis* induces relatively stronger immune reactions and pathologic changes in NMRI mice than newly derived isolates of both *E. ferrisi* and *E. falciformis*.

Between the two *E. falciformis* isolates BayerHaberkorn1970 and Brandenburg88 we observed only slight differences in the length of the pre-patent period (time until oocyst shedding, starting at 6 *vs.* 7 dpi). These results are in agreement with previous reports from the same host (NMRI mice) and the BayerHaberkorn1970 isolate (Stange et al., 2012; Schmid et al., 2012, 2014; Ehret et al., 2017). The pre-patent period for the wild derived *E. falciformis* isolate (7 dpi) corresponds to that reported for the parasite isolate *E. falciformis* var praghensis (Mesfin et al., 1978; Kasai et al., 1991), but Mahrt and Shi (1988) and Schito et al. (1996) demonstrated slightly longer pre-patent periods (7 or 8 dpi) also in other *E. falciformis* infections.

The output of oocysts in our study (for all isolates) was similar or only slightly lower than in previous reports (Schmid et al., 2014; Ehret et al., 2017). Our observation regarding the lifecycle progression of *E. ferrisi*, agree with the initial description of the life cycle in *Mus musculus* (Ankrom et al., 1975). *E. ferrisi* is characterized by a short life cycle - especially when compared to *E. falciformis* - with patency at 3 dpi. It is also noteworthy that the oocyst output of this species did not differ significantly when compared to that of both *E. falciformis* isolates (Figure 1). This data establishes *E. ferrisi* Brandenburg64 as a parasite isolate with low pathogenicity and high parasite reproduction. It invites comparisons of host-parasite interactions with more pathogenic isolates and could be an attractive alternative infection model with less impact on the host, i.e. allowing shorter experiments more compliant with welfare of host animals.

Pathogenicity assessed by maximal weight loss during infections with *E. falciformis* was more severe in the present study compared to previous experiments (Schmid et al., 2012; Stange et al., 2012; Ehret et al., 2017). A potential reason for such a relatively high pathogenicity could – as in any experimental *Eimeria* infection – be an underestimation when adjusting the number of sporulated oocysts for the inoculum. As observed by Haberkorn (1970) higher dosed inocula lead to enhanced pathology, while the extent and timing of oocyst shedding are not impacted. Importantly, we here developed a method to assess the intensity of intestinal stages relative to the number host cells (host-parasite ΔCt). This allows us to argue for the consistency of infectious doses: the tissue intensity attained by all three strains used in our experiment was not significantly different, and the tissue intensity of the two *E. falciformis* was very similar throughout the infection. We can conclude that we report results from a strong infection relative to previous studies, which was consistent between innocula of different strains.

This allows us to compare weight loss as an indication of pathogenicity between strains and to relate this to parasite reproduction: in *E. falciformis* infections maximal weight loss was observed at 9 dpi, while infections with *E. ferrisi* induced a significantly lower maximal weight loss at 5 dpi. These observations can be either due to fewer or faster cycles of asexual merogony of the parasite. Lines of poultry *Eimeria* with an abbreviated life cycle (so called “precocious lines”) show low oocyst output and are less pathogenic to their host (McDonald and Ballingall, 1983; Shirley and Bellatti, 1988; Shirley and Long, 1990; Shirley and Harvey, 2000). *E. ferrisi*, in contrast, attains substantial oocyst output, but shares low pathology with such precocious lines. This suggests that short phases of asexual expansion might be associated with low pathogenicity in *Eimeria* infections independent of total oocyst shedding. Parasite fitness for “naturally precocious” species of *Eimeria* might be high, while simultaneously host fitness is affected relatively little by infections. Short lifecycles of *Eimeria* might be associated with tolerance by the host.

Integrating weight loss dynamics with parasite lifecycle progression and comparing the two isolates of *E. falciformis* and that of *E. ferrisi* shows that *E. ferrisi* induces most weight loss before the peak of its oocyst shedding, while both *E. falciformis* isolates impact the host after the peak of their oocyst shedding (Figure 2). These differences suggest that mechanisms underlying pathogenesis might be different between the two parasite species. Histology indicates that weight loss coincides with immune cell influx in *E. falciformi*s infections (Figure 5). This influx differed slightly in timing starting at 5 dpi in the *E. falciformis* laboratory isolate and 7 dpi in the wild derived isolate. Such influx of immune cells into the tissue is an immuno-pathological reaction (Stange et al., 2012), which might cause damage to the host. As an alternative or additional cause of pathogenicity, sexual reproduction of *E. falciformis* might directly cause an exhaust of epithelial cell which burst when oocysts are released into the lumen (Kasai et al., 1991). Infections with *E. ferrisi* were characterised by yet lower immune cell infiltration and weight loss coincided with the peak abundance of endogenous stages at 3 dpi (Figure 2), indicating that parasite proliferation causes disease in host infected with this species.

Inflammatory cellular infiltrations within the mucosa during experimental infections with *Eimeria* have been observed in many host species including mice (Mesfin et al., 1978; Rose et al., 1992; Laurent et al., 2001; Gadde et al., 2009; Muñoz-Caro et al., 2016). Schmid et al. (2014) demonstrated by immunohistochemical analyses that *E. falciformis* infection in the caecum of NMRI mice leads to tissue infiltration with lymphocytes and macrophages. These changes are accompanied by elevated expression of *Infγ* and the production of the major chemokines CxC subfamily at the site of infection. Inflammatory infiltrates were also slightly more prominent in our experiment in the laboratory isolate *E. falciformis* BayerHaberkorn1970 than in the closely related wild derived *E. falciformis* Brandenburg88 isolate. We thus asked whether systemic immune response differs between infections and studied this based on mRNA expression in the spleen. In spleens of mice infected with the laboratory isolate of *E. falciformis* we observed elevated mRNA levels of the pro-inflammatory Th1 cytokine *Il12*. The *Il12*/*Ifnγ* axis is crucial for the activation of cellular immune responses against intracellular parasites including *Eimeria* (Ovington and Smith, 1992; Rose et al., 1992; Lowenthal et al., 1997; Lillehoj, 1998; Chow et al., 2011; Schmid et al., 2014; Ehret et al., 2017). We did not detect *Ifnγ* significantly unregulated itself, as an elevated expression was only observed early after infection (at 3 dpi). We, however, observed significantly increased expression of the anti-inflammatory Th1 cytokines *Il10* and *Tgfβ*. IL10 could counteract IFNγ and is also expressed in the spleen of *Eimeria*-infected chicken (Rothwell et al., 2000). Il10 expression in the spleen could be indicative for an attempt to balance inflammation during infection. A failure to establish this inflammatory balance can lead to pronounced inflammation and immunopathology (Inagaki-Ohara et al., 2006). In addition, we observed significantly elevated mRNA expression of *Stat6* and the major regulatory chemokine *CxCL9*, which can be induced downstream of INFγ and are involved in recruitment and activation of effector T lymphocytes in the spleen as well as in non-lymphoid organs such as intestine (Schmid et al., 2014). Only a few studies have assessed systemic immune response via expression of cytokines in the spleen during *Eimeria* infections. Steinfelder et al. (2005) showed that T-cell proliferated in mesenteric lymph nodes of mice during a drug abbreviated infection with *E. falciformis.* Nevertheless splenocytes released IFNγ and IL4 and likely contribute to the development of a systemic humoral response. In strongly infected mice this could lead to a systematic immunopathology. *E. tenella* antigen has been shown to induce IFN release of splenocytes in immunized chickens (Prowse and Pallister, 1989). Similarly, Byrnes et al., 1993 illustrated that splenic macrophages can produce IL1 and TNFα during the primary infection of *E. tenella* and *E. maxima*. The expression of Toll-like receptors (TLR3, TLR15), signal adaptor (MyD88) (Zhou et al., 2014) and IFNy (Rothwell et al., 2000) has been detected in the spleen of chickens as a response to infection with *E. tenella*. Taken together our data indicate a systemic immune response against *E. falciformis* BayerHaberkorn detectable in the spleen based on elevated mRNA levels of cytokines previously associated with in *Eimeria* infections.

Infection with the laboratory isolate of *E. falciformis* lead to significantly higher expression levels of cytokines in the spleen compared to both wild-derived isolates. In contrast, the wild derived isolates of *E. falciformis* and *E. ferrisi* did not induce significant expression changes over control levels in our mixed effect model analysis.

In spite of the overall non-significant changes, stints of elevated mRNA expression (failing to be significant in an analysis over different time points) seem plausible for *Il6* in the spleen during infections with all three *Eimeria* isolates. IL6 is expressed during the initial stage of inflammation at the site of infection, where it has a role in stimulating the intestinal epithelial proliferation and repair after injury (Kuhn et al., 2014). It can be transported as a protein through the bloodstream to the liver and spleen (Heinrich et al., 1990), where it promotes specific differentiation of naïve CD4+ T cells (Rincón et al., 1997). In experimental infections of mice with *E. falciformis* marked induction of *Il6* transcription between 5 and 7 dpi has been reported at the site of infection (Ehret et al., 2017). Enhanced expression in the spleen might suggest that the immune modulatory role of IL6 in the spleen during *Eimeria* infections could be augmented by elevated mRNA expression within this organ.

Similarly, *Tgfβ* mRNA expression levels seemed elevated in the spleen early during infection with *E. ferrisi* (3 and 5 dpi) and late in infections with the *E. falciformis* laboratory isolate (7 and 9 dpi; only in the latter significantly though). The simultaneous elevation of *Il6* expression levels, may indicate the involvement of a Th17 pathway to control the infection events. *Tgfβ* and *Il6* play crucial roles in the induction of IL17 expression from naïve CD4+ T cells of mouse (Sehrawat and Rouse, 2017; Korn et al., 2009). IL17 in turn contributes to both immunopathology and parasite restriction during infection with *E. falciformis* (Stange, 2013). These exceptions only underline the fact that a systemic immune response was hardly detectable, using gene expression in the spleen, in infections with wild derived *Eimeria* isolates.

The apparent differences in immune response of the wild derived and the laboratory isolate of *E. falciformis* invite speculation about their origin. Unfortunately, we do not know the infection phenotype (pathogenicity and induced immune reactions) of *E. falciformis* BayerHaberkorn1970 directly after its isolation. It is plausible, however, that the pathology before serial passaging resembled that observed for our wild derived isolate. The consequences of serial passaging could then be seen as the result of an accidental evolutionary experiment (Ebert, 1998).

Independent of the ultimate reasons for the difference in immunogenicity, we conclude that the infections with the laboratory isolate *E. falciformis* might not be representative for parasite-host interaction in their original ecological and evolutionary context. *Eimeria* is one of the most relevant parasites for wildlife immunology and, for representative infection experiments, we recommend to isolate strains of the parasite from the natural system with minimal prior passaging.

## 3. Material and Methods

### 3.1. Isolation of E. falciformis Brandenburg88 and E. ferrisi Brandenburg64

The pure inocula of wild derived isolates of *E. falciformis* Brandenburg88 and *E. ferrisi* Brandenburg64 were produced by a single passage through NMRI mice preceding the reported experiment. Briefly, sporulated oocysts of *Eimeria* for each isolate were recovered from faecal samples obtained by trapping individual wild house mice in the federal state of Brandenburg (Germany). The last part of the full name of novel isolates corresponds to a running number used for mice caught during field sampling. The mouse yielding the isolate *E. falciformis* Brandenburg88 had been caught at 13.84 52.2678 DD, the mouse yielding *E. ferrsi* Brandenburg64 at 13.4642 52.4164 DD.

Isolates were identified through amplification and sequence comparison of three genetic markers (small ribosomal subunit 18S, Cytochrome c Oxidase I COI and Open Reading Frame 470 ORF470) using primers previously reported (Zhao and Duszynski, 2001; Kvičerová et al., 2008; Ogedengbe et al., 2011). Sequences were generated by LGC (Berlin) based on forward and reverse primes and resulting forward and reverse reads were merged to a consensus sequence. Sequences from *E. falciformis* Brandenburg88 (MH751942, MH755305, MH755336) and *E. ferrisi* Brandenburg64 (MH751927, MH777469, MH755326) were deposited in the NCBI Genbank.

From flotations of these faecal samples 300 and 600 oocysts for *E. ferrisi* Brandenburg64 and *E. falciformis* Brandenburg88, respectively, were inoculated into 16 weeks old female NMRI mice. All mice were reared individually in wire cages in isolation cabinets and provided with food and water *ad libitum*. The faeces from those mice were collected daily. Oocysts in faeces were harvested using flotation (see below). They were then placed in 2% potassium dichromate and incubated at 25 °C for 4 days to permit oocyst sporulation. Sporulated oocysts were examined repeatedly under a light microscope and were then stored at 4 °C for about 1 month prior to use. The inoculum of *E. falciformis* BayerHaberkorn1970 was prepared simultaneously using the same protocol.

### 3.2. Infection protocol, oocyst counting and sample collection

60 female NMRI mice (10 to 12 weeks old) were randomly assigned to one of four groups, including a control group that was not inoculated. Oocyst concentrations were adjusted by counting the total number of oocysts in 10 μl, directly on a standard microscope slide. Using these inocula, 45 mice (15 per group) were inoculated via oral gavage with 0.1 ml of inoculum containing a single dose of 200 sporulated oocysts.

Faeces were collected daily, weighted and stored in 2% solution of potassium dichromate. For flotation the faecal material was homogenized, centrifuged at 3175g and the pellet was washed with distilled water. Oocysts were recovered from the sediment by 2 successive flotations in saturated NaCl solution each followed by washing. After the last washing 2 ml PBS were added, the pellet was suspended and 10 μl of the solution were loaded into a “Neubauer-improved chamber”. Oocysts were counted in eight grid squares. Then the number of oocysts per gram faeces was obtained according to the (0.1 μl) volume of one grid square: *Concentration (oocyst/g) = total #of oocyst / #squares counted * 10.000 ml-1 * 2ml / g (faeces).*

During the 11 days of the experiment the body weight of mice was recorded. From each group three mice were sacrificed on 3, 5, 7, 9, and 11 dpi. Immediately after death the viscera were exonerated and spleen and caeca removed. Caecal contents were gently removed with physiological NaCl solution and the tissue was cut longitudinally into two pieces. One piece was transferred into a 1.5 ml tube containing RNAlater® (Life Technologies; Carlsbad, USA) and stored for 4 h at 4°C before being transferred to −20 °C. The second piece of caecum tissue was fixed in 4% formalin and stored at room temperature for histological examination.

### 3.3. Quantification of *Eimeria* load in caecum tissue

Frozen caecum tissue was manually homogenized by grinding in liquid nitrogen. Genomic DNA was extracted using innu PREP DNA Mini Kit® (Analytik Jena; Jena, Germany) according to the manufacturer’s protocol incorporating proteinase K digestion. Purified DNA was stored at −20°C.

A Mitochondrial COI fragment of *Eimeria* spp. was amplified using primers Eim-COI-forward 5’TGTCTATTCACTTGGGCTATTGT3’ and Eim-COI-reverse 5’GGATCACCGTTAAATGAGGCA 3’. Host genomic DNA was amplified using a primer pair targeting the *Mus*-*cdc42* gene: *Mus*-*cdc42-* forward 5’CTCTCCTCCCCTCTGTCTTG3’ and *Mus*-*cdc42-*reverse 5’TCCTTTTGGGTTGAGTTTCC3’.

DNA samples were added for qPCR to Multiplate™ 96-Well PCR plates (BioRad; Hercules, USA), with reactions performed in duplicate for each sample. Each plate also contained a non template control and a plate control sample (ddH2O). The qPCR mixture was prepared using the iQ™ SYBR® Green PCR Kit (Bio-Rad; Hercules, USA): 5 μl of 2X iQ™ SYBR® Green Master Mix, 0.3 μl of 20 μM forward and reverse primers, and 4 μl of 10 ng/μl template DNA. The thermal cycling protocol was set as follows: initial denaturation for 15 min at 95°C, followed by 40 cycles of 15 sec at 95°C, 30 sec annealing at 60°C for Eim-COI primer or 58°C for *Mus*-*cdc42*, and 30 sec at 68°C and measuring the fluorescence signal at the end of every step.

qPCR amplifications were performed using the Bio-Rad CFX96, Thermalcycler1000 system, which determined the cycle at threshold (Ct) fluorescence. To confirm the specificity of the assay a melting curve was generated after RT-PCR by adding a stepwise temperature increase from of 65.0°C to 95.0°C, with 0.5°C increment. After calculating mean Ct of technical replicates, the abundance of *Eimeria* relative to host DNA was estimated as the ΔCt between mouse and parasite DNA. This is equivalent to a log(2)-ratio between mouse host (*Mus*-*cdc42*) and *Eimeria* parasite (Eim-COI) DNA copies. The number of copies was calculated by taking the antilog for some examples in the text.

### 3.4. RNA extractions and reverse transcription

Frozen spleen tissue was homogenized by grinding in liquid nitrogen. Total RNA was isolated using the PureLink™ RNA Mini Kit (Thermo Fisher Scientific; Waltham, USA). Briefly, frozen homogenized sample was transferred with a sterile scalpel blade into tubes with 1ml lysis solution with 1% 2-Mercaptoethanol and 1.4 mm zirconium oxide beads (Peqlab GmbH, Erlangen, Germany). Samples were homogenized at room temperature using a Precellys® 24 tissue homogenizer (VWR; Radnor, USA) twice at 6,000 rpm for 20 sec interrupted by a 30 sec cooling break. Samples were centrifuged for 1 min at maximum speed (13,400 rpm) to eliminate foam. Supernatant was collected and mixed at a 1:1 ratio with 70% Ethanol. Afterwards, 600 μl of the mixture was added on a spin filter and centrifuged at 13,400 rpm for 30 sec. On-column DNase digestion was performed using PureLink DNase (Thermo Fisher Scientific, Waltham, USA) according to the manufacturer protocol. Column washing and elution was preformed as indicated by the manufacturer.

Synthesis of complementary DNA (cDNA) was performed using the RevertAid H Minus First Strand cDNA Synthesis Kit (Thermo Fisher Scientific, Waltham, USA) with engineered RevertAid™ H Minus M-MuLV Reverse Transcriptase (200 U/μl). Nuclease-free water and 2 μl Oligo (dT)18 primers (100 μM, 0.5 μg/μl) were added to 1 μg template RNA to a total volume of 22 μl. To denature potential secondary structures, the mixture was heated to 65°C for 5 min using the 2720 Thermal Cycler (Applied Biosystems; Foster City, USA) and rapidly cooled on ice afterwards to prevent renaturation. Reverse transcription was carried out adding the reverse transcriptase mix and incubating for 60 min at 42°C followed by a termination at 70°C for 10 min.

### 3.5. Gene expression quantification

We measured the mRNA expression levels of seven target genes: *CxCL9, Il10, Il12, Tgfβ, Stat6, Il6 and Infγ. Cdc42, Ppia* and *Ppip* were confirmed to be suitable as reference genes using 16 randomly selected cDNA samples in an analysis of mRNA expression stability (Axtner and Sommer, 2009; Weyrich et al., 2010) performed in qbase+ (Biogazelle; Zwijnaarde, Belgium) as implemented in the Bio-Rad CFX96 Thermalcycler1000. Respective primers are given in Table 3.

**Table 3:**
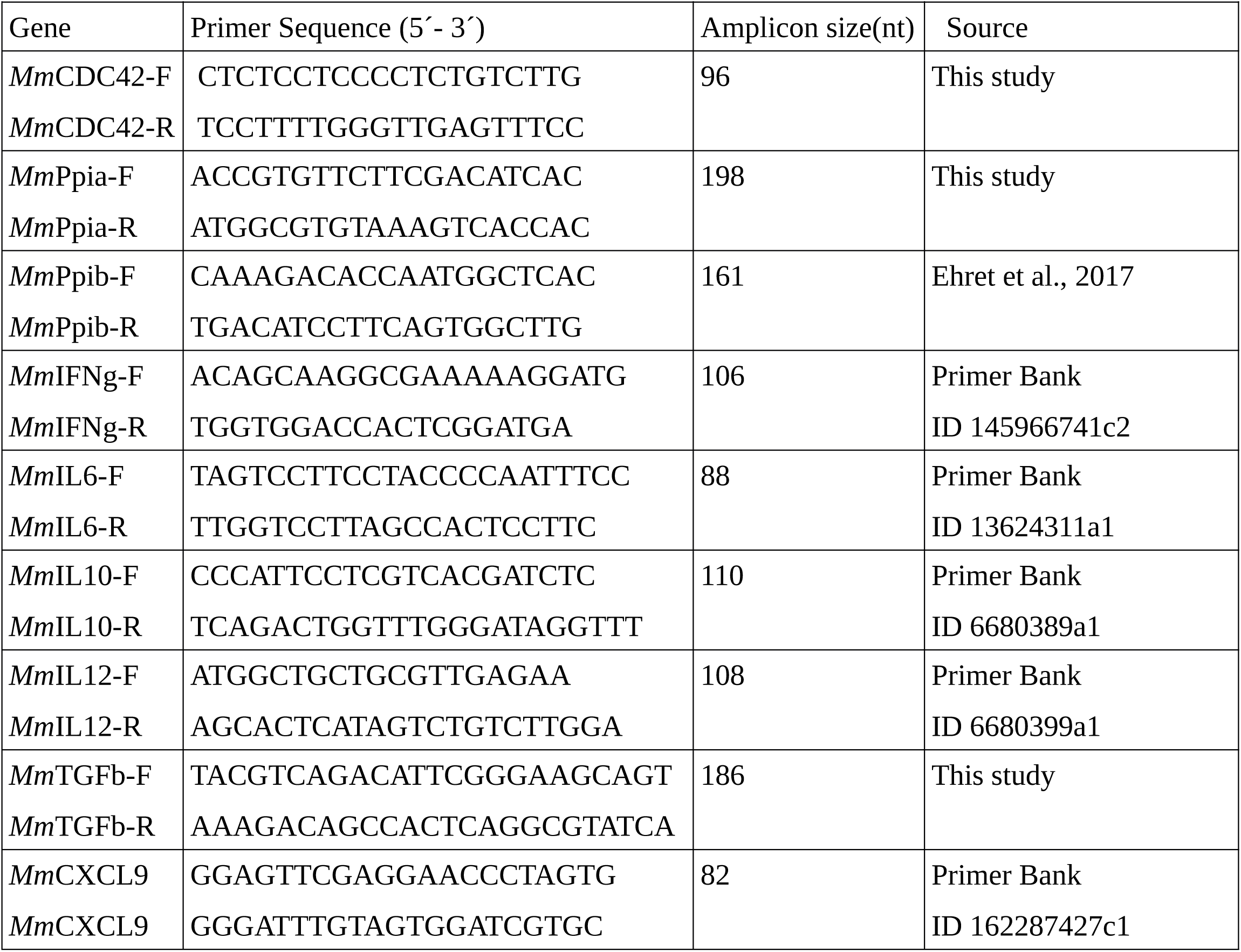
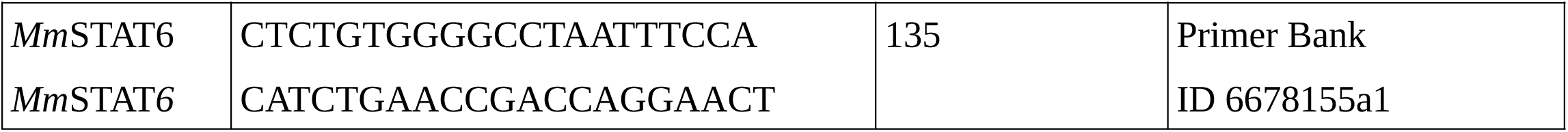
Primer-pairs used for qPCR analysis of gene expression. cDNA samples were split between PCR plates (Multiplate™ 96-Well, Bio-Rad; Hercules, USA) with reactions performed in duplicate for each sample. If the standard deviation of Ct values between duplicates was > 0.4, reactions for corresponding samples were repeated (as described in Weyrich et al., 2010). Each plate contained a non-template control sample.

The qPCR mixture of 10 μl was prepared using the iQ™ SYBR® Green PCR Kit (Bio-Rad): 5 μl of 2x iQ™ SYBR® Green Master Mix, 3 μl of 10 μM forward and reverse primers, and 4 μl of 10ng/μl template cDNA. qPCR amplifications were performed using Bio-Rad CFX96, Thermalcycler1000 system as follows: initial denaturation for 15 min at 95°C, followed by 40 cycles of 15 sec at 95 °C, 30 sec at 60 °C and 30 sec at 68 °C with a measuring of the fluorescence signal at the end of every step. The cycle of quantification (Ct) was determined by the amplification plot in CFX96-Bio-Rad software. Finally, a melting curve was generated to confirm the specificity of the reaction by adding a cycle of 65°C to 95°C in 0.5°C increments.

Normalization factors (NF) were calculated using the geometric mean of expression values for the three reference genes (Vandesompele et al., 2002). Relative expression values for each tested sample and each gene of interest were then calculated using the ΔCt method, adjusted for the amplification efficiencies of each primer pair and standardized against the NF of each sample.

### 3.6. Histological examination and scoring

Formalin fixed samples from the mid-part of the caeca were embedded in paraffin and sectioned with 4 μm thickness. Tissue slides were stained with hematoxylin and eosin and were examined at 100-200- and 400-times magnification by light microscopy. The extent and nature of leukocyte infiltration in the intestinal wall was assessed based on the morphological characteristic of each cell type. A numerical score was assigned with 0 representing no leukocyte infiltration and 1, 2, and 3 mild, moderate, or severe infiltration, respectively.

### 3.7. Statistical analyses and visualisation

All statistical analyses and visualisations were performed in R (R Development Core Team, 2008). The non-parametric Mann-Whitney U-test was used to assess differences in the distributions of weight loss or oocyst shedding. Linear mixed effect models (function “lmer” of the package lme4) were used to test for differences in gene expression. For each gene (as response variable) these models used the infecting *Eimeria* isolate as only fixed effect and the time of infection (dpi) as a random intercept. Similarly, linear mixed effect models for leukocyte infiltration score (as response variable) were used with infection isolate as fixed effect and dpi as a random intercept. For visualisations the package ggplot2 was used, including the default “loess” smoother as indicated in figure legends.

## Ethics statement

All animal procedures in this investigation were performed according to the German Animal Protection Laws as directed and approved by the overseeing authority Landesamt für Gesundheit und Soziales (Berlin, Germany) under permit number H0098/04.

## Acknowledgements

The authors thank Deborah Dymke and Jenny Jost for help during the infection experiment, Anke Schmidt and Doris Krumnow for excellent technical assistance in qPCR and histology, respectively.

